# Ripples at Edges of Blooming Lilies and Torn Plastic Sheets

**DOI:** 10.1101/2020.08.05.235176

**Authors:** Thomas Portet, Peter N. Holmes, Sarah L. Keller

## Abstract

Ripples arise at edges of petals of blooming *Lilium casablanca* flowers and at edges of torn plastic sheets. In both systems, ripples are a consequence of excess length along the edge of a sheet. Through the use of both time-lapse videos of blooming lilies and still images of torn plastic sheets, we find that ripples in both systems are well-described by the scaling relationship 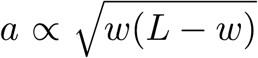, where *a* is amplitude, *w* is wavelength, and *L* is arc length. By approximating that the arc length is proportional to the wavelength, we recover a phenomenological relationship previously reported for self-similar ripple patterns, namely ⟨*a*⟩ ∝ ⟨*w*⟩. Our observations imply that a broad class of systems in which morphological changes are driven by excess length along an edge will produce ripples described by 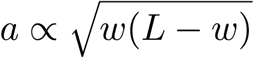.

**Significance Statement:** Early in the blooming process of *Lilium casablanca* flowers, large ripples appear in the edges of their petals. As blooming progresses, smaller ripples arise on top of the original ones. All ripples are characterized by three variables, an amplitude, a wavelength, and an arc length. Here, we derive an equation that relates these variables. To test the equation, we collect movies of blooming lilies. We find that the equation quantitatively describes single ripples in the petals. The equation is general, so it applies to all systems in which ripples arise by the same mechanism. To illustrate this point, we show that the equation holds for single ripples that appear at the edges of torn plastic sheets.

## Main Text

Morphological changes in flowers elegantly connect the abstract beauty of mathematics and the tangible beauty of nature. Two recent studies inspired our current investigation of lilies. In the first, the authors applied a growth hormone to the edge of flat plant leaves to create ripples (*1*). In the second, the authors combined surgical manipulations of *L. casablanca* lilies, numerical simulations, and exact calculations to argue that lily blooming is driven primarily by edge growth (*2*). In this mechanism, growth along edges of lily tepals (colloquially termed petals) is enhanced with respect to tepal faces, such that excess length accumulates along edges. Blooming of lilies from closed buds to six-pointed stars (Fig. 1) entails curvature reversal of tepals from cup shapes to saddle shapes. Edge growth is not the only possible mechanism of blooming. Historically, lily blooming has also been attributed to enhanced expansion of cells on the adaxial (interior) face of tepals with respect to cells on the abaxial (exterior) face. Bieleski *et al*. assert that any differential expansion of this type must play a minor role because it follows (rather than precedes) the onset of blooming (*3*). Changes in the angle and curvature of the midrib—the tepal’s woody spine—also contribute to blooming (*2, 3*). In other types of petals, differential rates of growth are less important than directional growth, as in *Antirrhinum* (snapdragon) (*15*), and cell-shape anisotropy, as in *Aquilegia* (columbine) (*16*).

**Figure 1:**
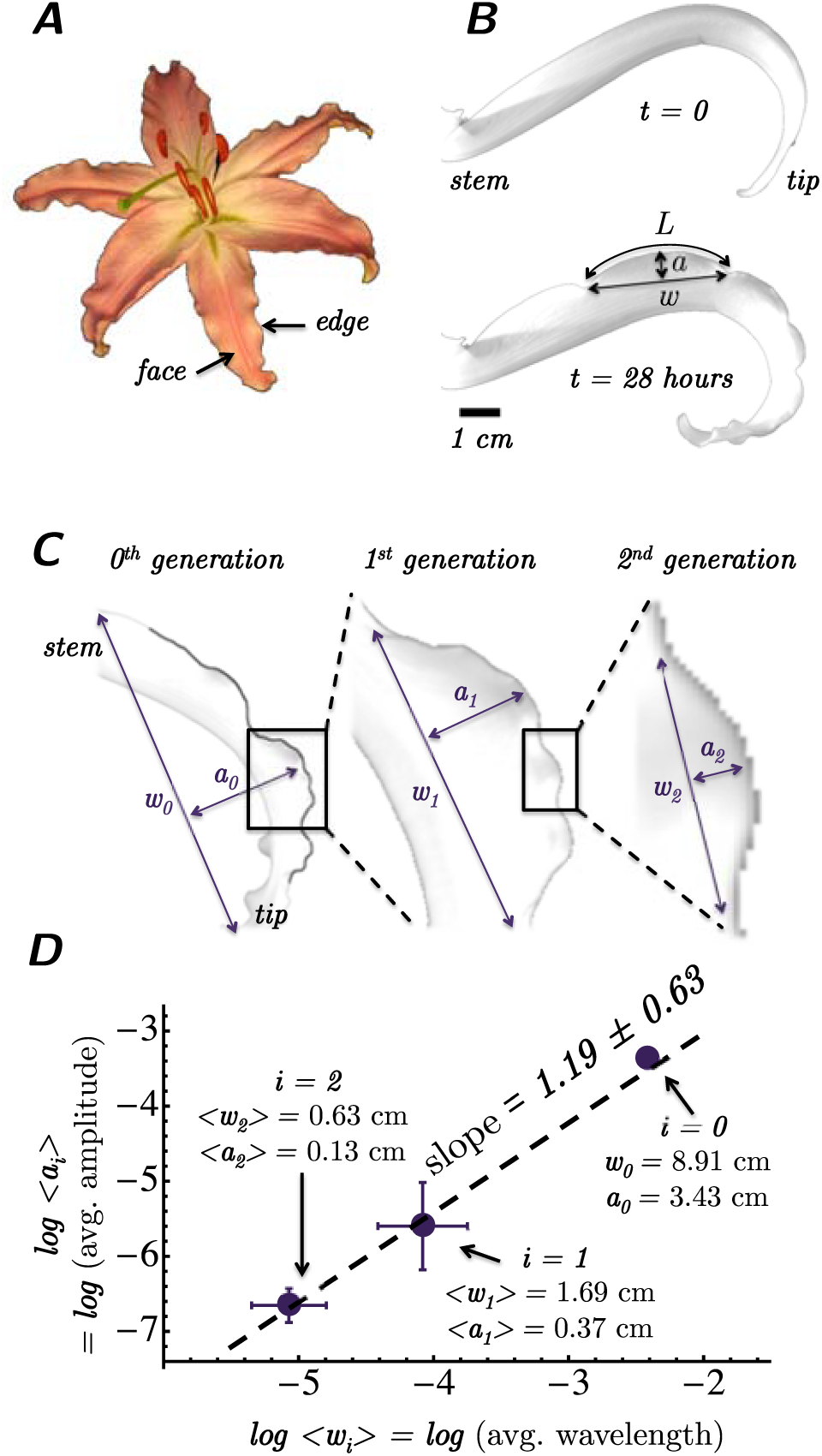
Self-similar ripples in lilies. **A:** Top view of an open stargazer lily with three inner and three outer tepals. **B:** Side view of an outer tepal of *Lilium casablanca* before (top) and after (bottom) edge rippling, with amplitude *a*, wavelength *w*, and arc length *L*. **C:** Examples of an amplitude *a*_*i*_ and wavelength *w*_*i*_ for ripples for each generation *i*. The leftmost image is outlined in black. **D:** Log-log plot of average final ripple amplitude ⟨*a*_*i*_⟩ *vs*. average final wavelength ⟨*w*_*i*_⟩ for each generation *i*. Fig. S1 labels all of the ripples analyzed in panel D. Error bars are standard deviations, and numbers of measurements are *N*_0_ = 1, *N*_1_ = 5, *N*_2_ = 6. Lily sizes limit observable length scales to one order of magnitude.

Ripples are observed in a wide class of systems that accommodate excess length along an edge. Simulations of elastic ribbons with high rates of edge growth result in rippled morphologies (*4*). Similarly, polymer disks undergoing nonuniform swelling buckle at their rippled edges, as do annular thin strips undergoing nonuniform stresses (*5–8*). In more quotidian examples, knitters create ripples by inserting stitches at edges, and ripples form at the new edges of a torn plastic sheet (*1, 9, 10*). Our goal is to quantitatively explain the shape of a single ripple in a sheet undergoing edge growth. Here we derive a simple scaling law of geometric origin to describe a ripple in terms of its amplitude *a*, wavelength *w*, and arc length *L*, and we find strong agreement between our scaling relationship and experimental data for single ripples at edges of lily tepals and torn plastic sheets.

Ripples in lily tepals appear singly or in multiple generations (Fig. 2). Fig. 1C shows a self-similar ripple pattern in a lily tepal in which a large first-generation ripple is decorated with smaller second-generation ripples. These patterns are reminiscent of self-similar patterns previously observed in edges of torn plastic sheets by Sharon *et al*. (*9*). In their 2002 paper, the authors demonstrated self-similarity of six generations of superimposed ripples in the buckled edge of a torn plastic sheet by fitting the relationship

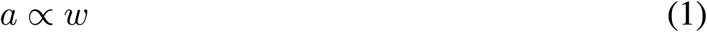

to data spanning 2.5 orders of magnitude in *w*, as summarized in Fig. 2 (*9*). However, we find that self-similar ripples at the edges of lily tepals are described by Eq. (1) only when all amplitudes and wavelengths within each generation are averaged (Fig. 1C-D). Additional reasons that the relationship *a* ∝ *w* is not entirely satisfying are that (i) the relationship is phenomenological and lacks physical justification, (ii) there is ambiguity in how to define *a* and *w* when multiple generations of ripples are present (Fig. 2), and (iii) fitting Fig. 1D to a 90% confidence interval results in a large (*≈* 50%) measurement uncertainty in the slope of *log*⟨*w*⟩ *vs. log*⟨*a*⟩, where the slope is 1.19±0.63.

**Figure 2:**
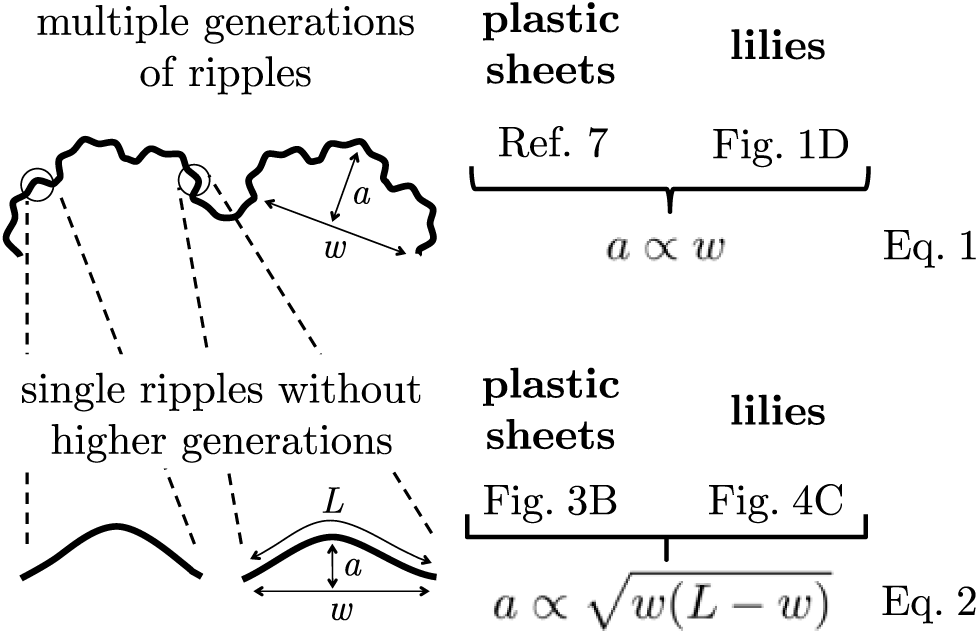
Summary figure. Generations of ripples at the edges of lily tepals and plastic sheets follow the phenomenological relationship *a* ∝ *w*. Single ripples are described well by the relationship 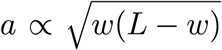, derived in the Supporting Material. The relationship *a* ∝ *w* is recovered by approximating *L* ∝ *w*.

We address the problems above by considering only single ripples, which we define as having no higher-order generation ripples superimposed upon them. In the Methods of the Supporting Material, we show that the amplitude *a*, wavelength *w*, and arc length *L* of single ripples should be related by:

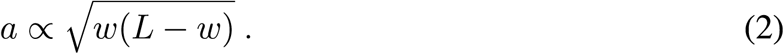

In Fig. 3, we show that this relationship holds for single, 6th generation ripples at the edge of the same plastic sheet studied by Sharon *et al*. (*9*), whereas the previously reported phenomenological relationship of *a* ∝ *w* does not.

**Figure 3:**
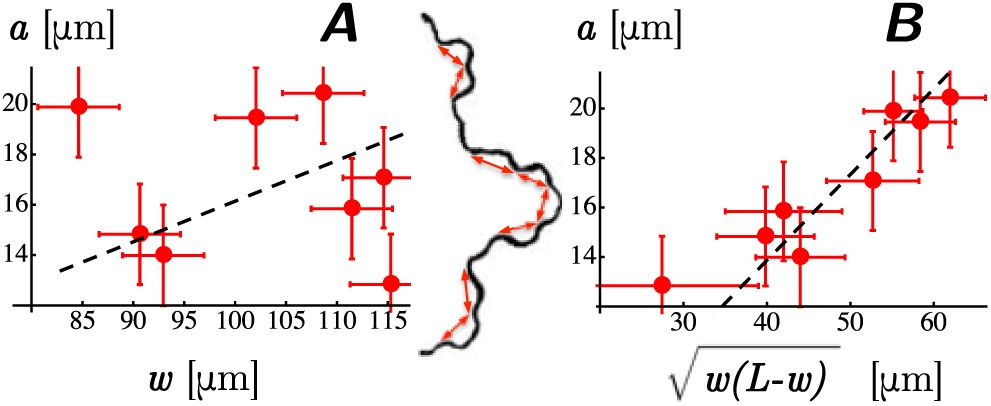
Tests of relationships between amplitude *a*, wavelength *w*, and arc length *L* for single ripples in plastic sheets. The image at the center shows single, 6th-generation ripples (*N* = 8) in a plastic sheet viewed from the edge (adapted from (*9*)). **A:** The shapes of these ripples are fit poorly by the dashed line of *a* ∝ *w*. **B:** The same ripple shapes are fit well by 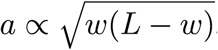, consistent with the prediction of Eq. (2).

Next, we collected videos of lilies blooming and rippling to show that single, 2nd generation ripples at edges of tepals are also well described by Eq. (2). *L. casablanca* specimens are ideal for video imaging because buds are large (∼ 10 cm), are widely available, and bloom over the course of approximately one day (*11*). We imaged tepals from the side in a time-lapse chamber (*12*). We find that Eq. (2) holds for single ripples in tepals of *L. casablanca* (Fig. 4), whereas all other scaling relationships fail in which amplitude *a* is proportional to a simple expression with units of length, namely *w, L*, 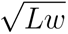, *L* + *w*, or *L* − *w* (Fig. S4). For clarity, Fig. 4 displays data for only single ripples, excludes ripples with amplitudes ≤ 0.1 cm, and does not show experimental uncertainties. Full data sets with experimental uncertainties appear in Fig. S5.

**Figure 4:**
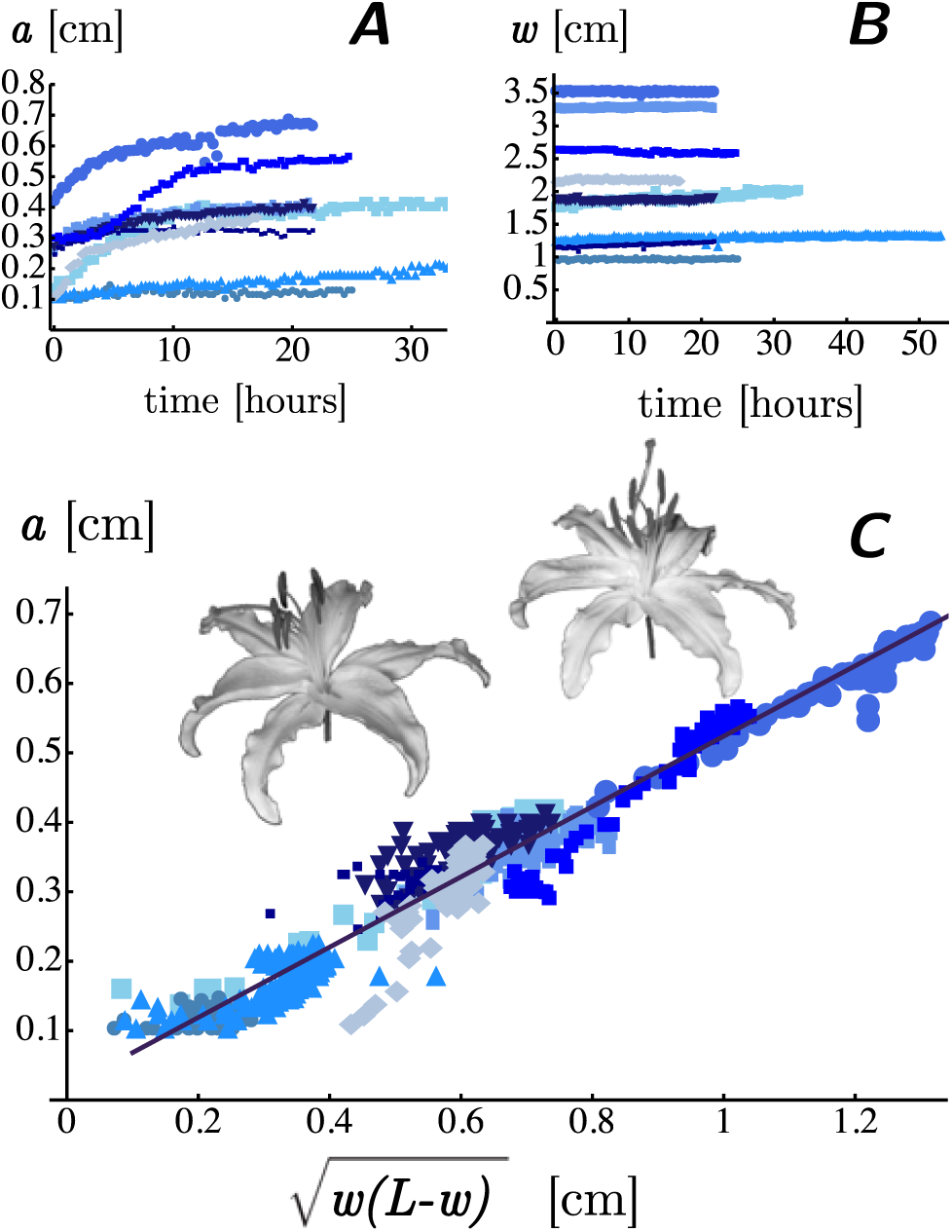
Relationships between amplitude *a*, wavelength *w*, and arc length *L* for nine first-generation ripples in lily tepals. Fig. S2 contains images of all ripples analyzed for this figure. A: Amplitude increases with time (as does arc length, see large-format graphs in Fig. S3). **B:** In contrast, wavelength *w* remains approximately constant over time. **C:** Within uncertainty, all data collapse onto a straight line when plotted as 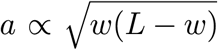. For clarity, this figure displays data for only single ripples with no superimposed higher generation ripples, excludes ripples with amplitudes ≤ 0.1 cm, and does not show experimental uncertainties. Full data sets with experimental uncertainties appear in Fig. S5.

Above, we have shown that the scaling relationship 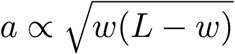 applies to a large population of single ripples at edges of both torn plastic sheets (Fig. 3B) and lily tepals (Fig. 4C). Next, we quantitatively reconcile this relationship with the phenomenological relationship *a* ∝ *w* reported for multiple generations of superimposed ripples (*9*). First, we approximate that *L* ∝ *w*. This proportionality captures the simplest possible relationship between arc length and wavelength — in practice, *L* is difficult to measure in a self-similar rippled edge. Inserting *L* ∝ *w* (where *L > w*) in Eq. (2), yields *a* ∝ *w*. Next, by defining the proportionality factors *γ* and *β* such that *L* = *γw* and 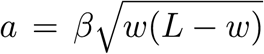, we find 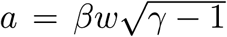. This equation successfully yields values that are consistent with previously published results for ripples in torn plastic sheets. To find *β*, we fit a straight line to the data in Fig. 3B, yielding a slope of *β* = 0.35 ± 0.02. To find *γ*, we fit a straight line to the data for *L* vs. *w* in Fig. S6, which yields *γ* = 1.22 ± 0.08. Combining *β* and *γ* gives 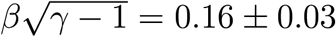, in agreement with the published value of 0.15 (*9*). Our simple, analytical approach produces a scaling relationship that accurately describes single ripples observed in sheets with edge growth. A previous, complementary computational approach considered a thin, elastic strip along a rippled edge. If disconnected from the larger sheet, a strip of this type would curl into a flat annulus. Given particular physical values characterizing each individual strip, numerical minimizations of the strip’s energy result in rippled solutions (*7, 8*).

Our analysis above shows how biology and physics can work in tandem to create lily petals with delicate ripples along their edges: genes regulate the rate of edge growth (*14*), and stresses within each tepal convert this growth into wavy edges. Within two-dimensional sheets, preferential wavelengths emerge from a competition between stretching and bending – the former favors short wavelengths, and the latter favors longer ones (*13*). As a tepal edge grows, small, first generation ripples appear at sites separated by a distance *w* that presumably arises from minimization of the sum of stretching and bending energies. The value of *w* remains approximately constant. This result would be expected if a tepal’s elastic modulus does not change significantly over time. As the tepal’s edge grows, the ripple’s arc length *L* increases. The tepal reacts to the excess edge length by buckling, just as a sheet of paper buckles when compressed laterally. Hence, ripple amplitude *a* grows as *L* grows. The magnitude of this growth is given by Eq. (2) such that *a* scales as 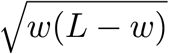. Identical ripples should arise in any material that undergoes similar stresses.

We have treated tepal ripples as a direct consequence of the same edge growth that drives lily blooming. In other words, any flower that exhibits ripples along an edge, whether as part of a lily tepal or a daffodil trumpet (*1*), does not necessarily genetically encode rippling beyond driving a generalized edge growth that results in blooming. The single ripples that we investigate may occur on their own or in a series of generations. Creation of hierarchical morphologies through nonuniform growth rates is of central importance in a range of biological systems, including intestinal loops of the mammalian gut (*17*). Our results set no constraints on the method by which a lily tepal achieves edge growth; any biological mechanism is allowed. Once edge growth occurs, a balance of physical forces determines the final tepal shape.

Here, we considered rippling due to edge growth in the absence of external forces. For long kelp leaves, the magnitude of flow in the surrounding water influences whether growing blades adopt flat or rippled morphologies, and pushing on a blade’s edge can alter the wavelength and amplitude of ripples (*19*). Similarly, in lotus leaves, the wavelength of edge ripples and the overall leaf shape are influenced by whether the leaves rest on a water surface or not (*18*). Together, these studies illustrate ways that edge growth in biological and biomimetic systems can be harnessed to achieve morphologies that are tuned by environmental conditions.

## Supporting Material

Supporting Methods, 1 movie, and 6 figures are available.

## Author Contributions

T.P. and S.L.K. designed research and wrote the paper. T.P. collected data and derived equations. T.P. and P.N.H. analyzed data.

## Acknowledgements

We thank Jennifer Nemhauser and her group for access to and assistance with time-lapse imaging. Lilies were from a local florist or the kind gift of Dianne Carlson, Marie Davis, Elizabeth Harasek, and Beth Hammermeister. We thank L. Mahadevan for his UW seminar that inspired this study, and we thank Jonathan Litz for his critique of our manuscript. S.L.K. thanks the Whitely Center at UW’s Friday Harbor Laboratories for space and quiet to write. Research in the Keller Lab is funded by NSF MCB-0744852 and MCB-1925731. T.P. was funded by the Raymond and Beverly Sackler Foundation and the Fondation Bettencourt Schueller.

## Acknowledgements

We thank Jennifer Nemhauser and her group for access to and assistance with time-lapse imaging. Lilies were from a local florist or the kind gift of Dianne Carlson, Marie Davis, Elizabeth Harasek, and Beth Hammermeister. We thank L. Mahadevan for his UW seminar that inspired this study, and we thank Jonathan Litz for his critique of our manuscript. S.L.K. thanks the Whitely Center at UW’s Friday Harbor Laboratories for space and quiet to write. This research was funded by NSF MCB-0744852 and MCB-1925731, the Raymond and Beverly Sackler Foundation (to T.P.), and the Fondation Bettencourt Schueller (to T.P.).

## Supporting Materials

### Movie legend

The supporting movie shows the rippling of the tepal displayed in Fig. 1B of the main text. Each ripple edge is highlighted in a different color. Arrows indicate wavelengths and amplitudes. Each frame corresponds to 20 minutes.

### Methods

#### Derivation of Eq. (2)

We consider the simple case of a first generation ripple that has no second generation ripples superimposed on it. We model the ripple’s edge as a planar curve for which each point has coordinates (*x, f* (*x*)), with *x ∈* [−*w/*2, *w/*2]. We assume the ripple function *f* is smooth, is even, and satisfies boundary conditions of *f* (*w/*2) = *f* (−*w/*2) = 0, and of *f* (0) = *a*. The arc length *L* of the curve is, by definition, 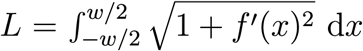, which is simplified via a Taylor expansion to 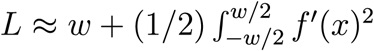 for a smooth function *f*. Because the function *f* is even, its expansion does not contain odd powers and is, to second order, *f* (*x*) *≈ a* + *f* ^*″*^ (0)*x*^2^*/*2. Using a difference quotient approximation, the second derivative at the origin is *f* ^*″*^ (0) *≈*−*E*(*a/w*^2^), with dimensionless factor *E*. Direct integration of *f* ^*′*2^ yields Eq. (2).

#### Videos and Measurement Uncertainty

Videos were collected of lilies blooming and rippling. Lilies of *L. casablanca* and *L. stargazer* varieties, which are Division VII oriental hybrids, were trimmed so only one outer tepal remained. The remaining tepal was imaged from the side at constant illumination of 60 *µ*mol*/*m^2^s and 20*°*C in a time-lapse chamber described previously by J. L. Stewart Lilley, C. W. Gee, I. Sairanen, K. Ljung, and J. L. Nemhauser, *Plant Physiol*. **160**, 2261 (2012). Image capture started a few hours after blooming commenced (before rippling) and stopped when ripples became static. Measurement uncertainties for *a, w* and *L* are equivalent and on the order of ±0.05 cm. Uncertainties arise mainly because tepal edges deviate from the camera’s focal plane due to flower motion or tepal shape. Smaller uncertainties arise from image analysis, which combines manual detection of ripple zeniths and nadirs with automated edge detection by ImageJ (National Institutes of Health, Bethesda MD).

### Figures S1 to S6

**Figure S1:**
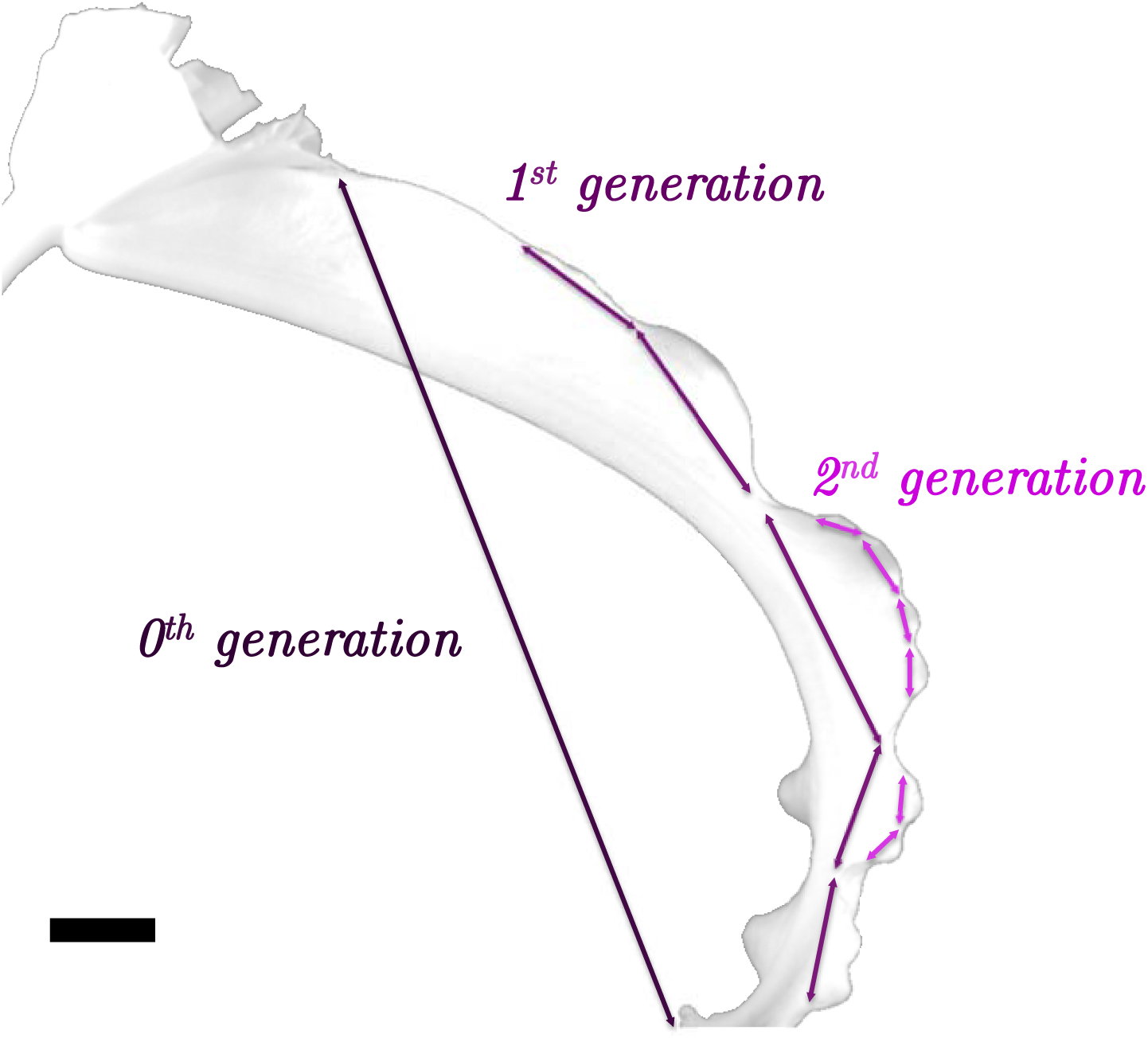
Index of ripples analyzed in Fig. 1. Photograph of *L. Casablanca* tepal ripples of generation 0, 1 and 2 used to obtain the plot of log(*a*) vs. log(*w*) in Fig. 1 of the main text. In the image, the stem of the lily is at the left, bases of five excised tepals are at the top, and the tip of the single remaining tepal is at the bottom. Scale bar: 1cm.

**Figure S2:**
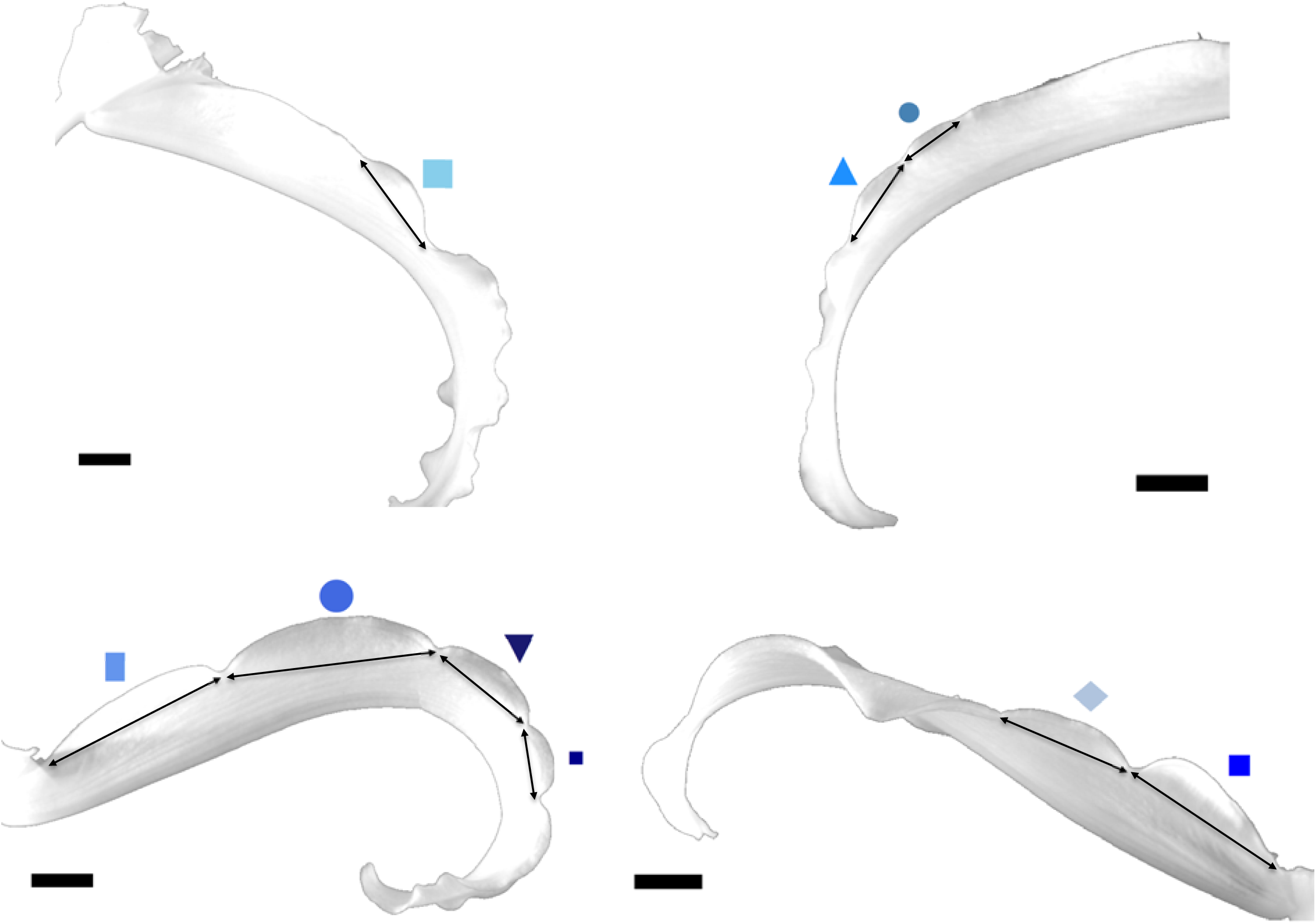
Index of ripples analyzed in Fig. 4. Photograph showing all simple, first generation lily tepal ripples evaluated. These ripples were chosen because they were smooth (*i*.*e*. without smaller, second generation ripples) and because their edges were straightforward to track automatically (*i*.*e*. their white edges were imaged against a dark background rather than against another part of the white tepal). Symbols correspond to those in Fig. 4 of the main text, and to those in Figs. S3, S4 and S5. In all cases, tepal stems are toward the figure edges. Scale bars are 1 cm.

**Figure S3:**
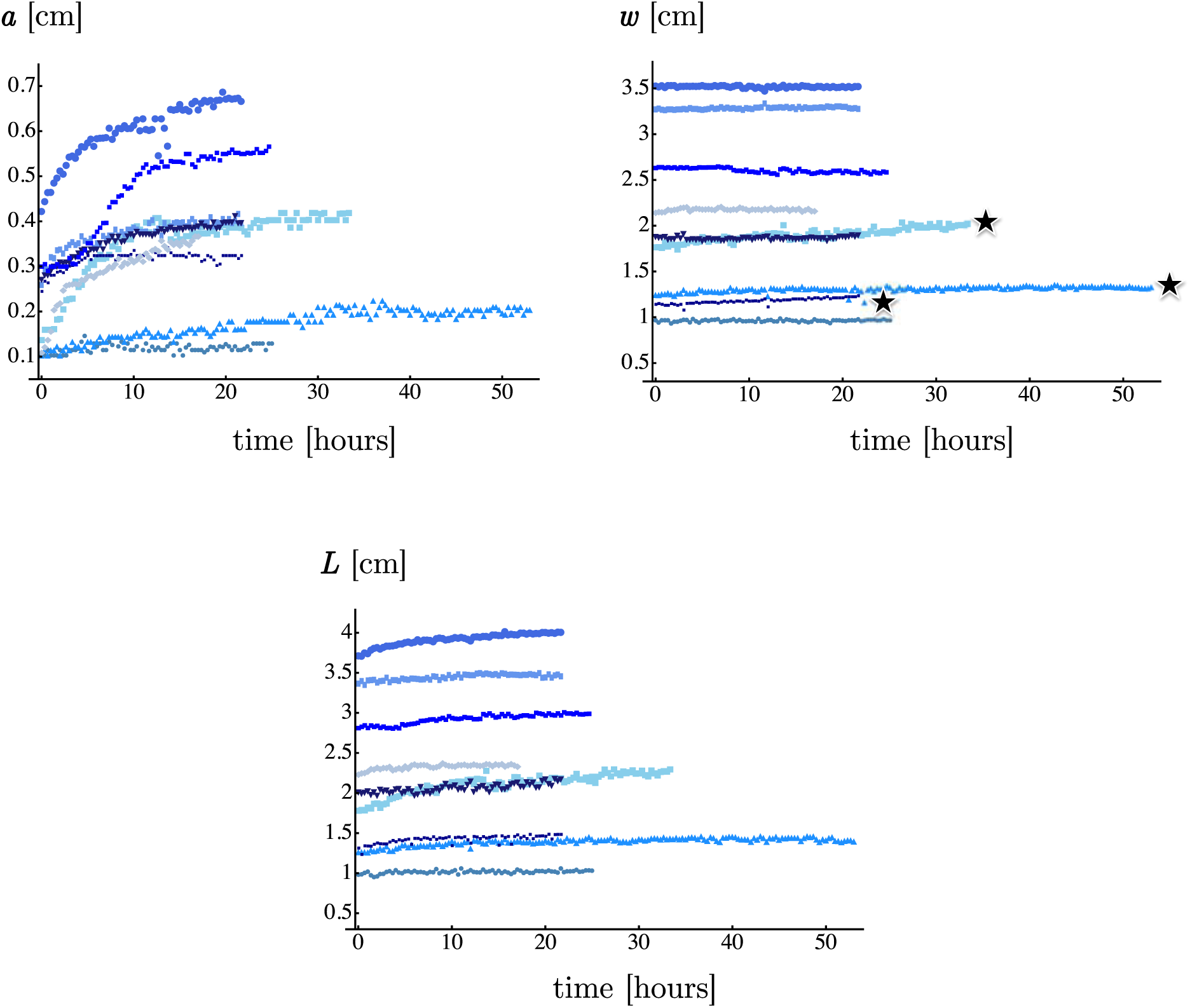
Evolution of shape descriptors. The scaling analysis presented in main text is restricted to simple, first generation ripples whose amplitudes are greater than 0.1 cm, well above our detection limit. (Data for smaller ripples are presented in Fig. S5. Here we show the evolution of three shape descriptors (amplitude *a*, wavelength *w*, and arc length *L*) throughout time for all simple ripples analyzed, namely those shown in Fig. S2. Here, the term “simple” means that first generation ripples were analyzed only if they contained no second generation ripples, and when second generation ripples were analyzed, the underlying first generation ripples were not analyzed. The top left panel of this figure shows that amplitudes increase with time and then plateau at final values. The top right panel shows that wavelengths remain approximately constant in time. This same attribute can be observed qualitatively in the supporting movie. The slight increase in *w* through time seen in the three data sets marked with stars (out of nine) is artifactual and due to gradual rotation of one of the tepals toward the camera during imaging. The bottom panel shows that for all ripples, arc length *L* increases and then plateaus.

**Figure S4:**
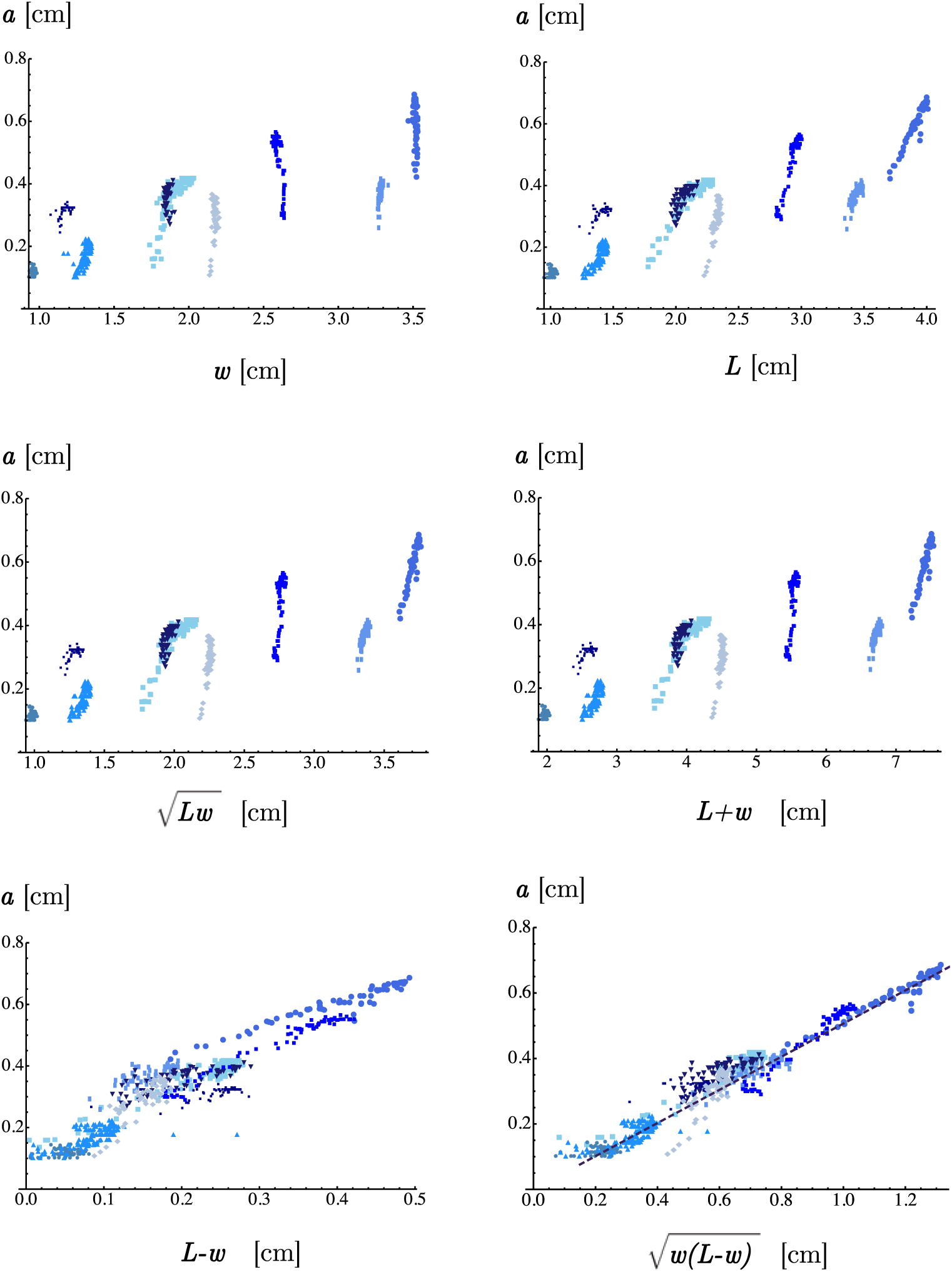
Examples of incorrect scaling relationships. In the main text, we derive the following simple scaling relationship between the three shape descriptors: 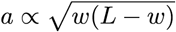. Fig. 4C in the manuscript is reproduced at the bottom right of this figure and shows that our observations are in excellent agreement with the relationship above. Namely, data collapse onto a straight line in ascatter plot of *a* vs. 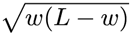. The purpose of this figure is to illustrate that collapse of the data by *a* vs. 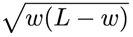 is superior to collapse of the data by any other simple relationship between *a* and *d*, where *d* is a distance constructed from various manipulations of *w* and *L*. For example, the top left panel shows *a* vs. *w*. As discussed in the main text, the scaling relationship *a* ∝ *w* does not apply to lily tepal edges when considering a single generation of ripples. The next three plots yield similarly poor collapses for *a* vs. *d* where *d* = *L* (top right), 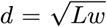 (middle left), and *d* = *L* + *w* (middle right). A better collapse is found for *a* vs. *L* − *w* (bottom left). The best collapse is found for *a vs*. 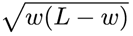 (bottom right).

**Figure S5:**
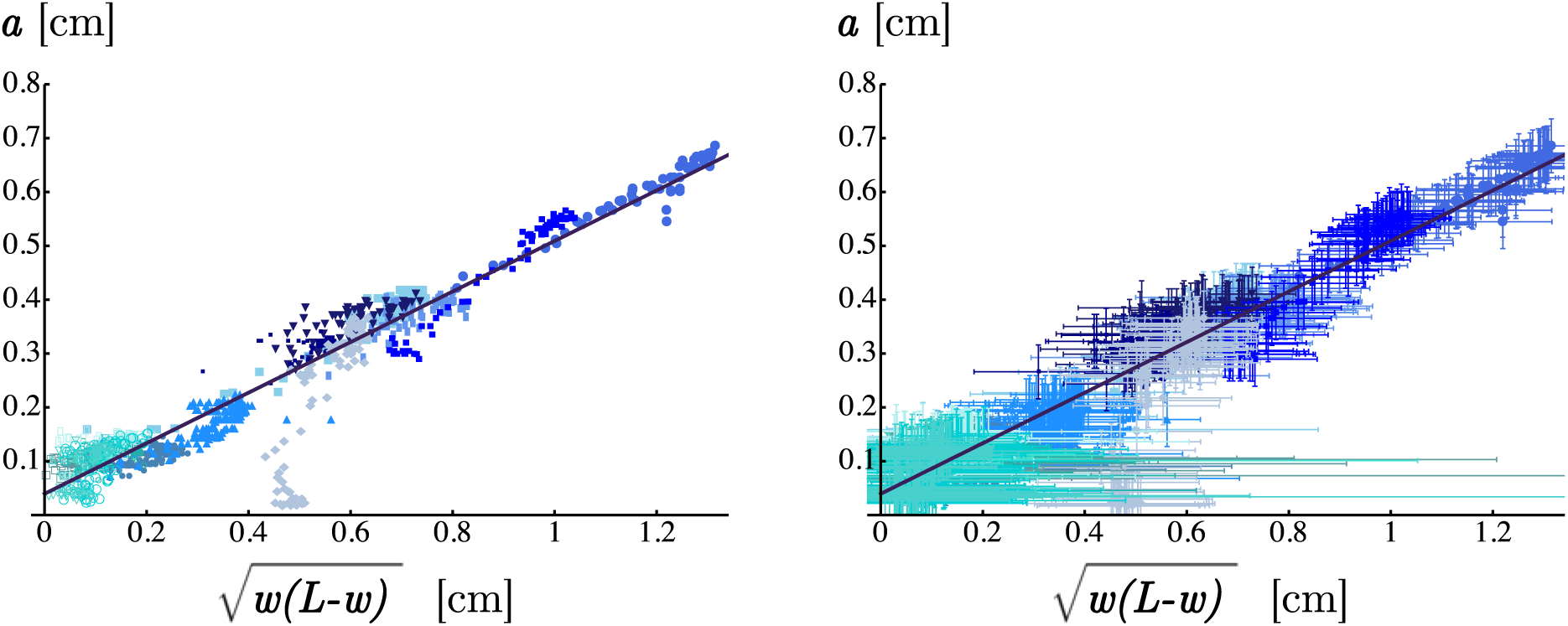
Data for smaller ripples. Data presented in Fig. 4C of the main text were restricted to ripples whose amplitudes were greater than 0.1 cm, well above the detection limit. This cutoff excludes second generation ripples and small first generation ripples. Here we present corresponding figures that include all simple ripples analyzed, including those with amplitudes smaller than 0.1 cm. Although the amplitudes of small ripples approach our measurement uncertainty, we find that all six of the second generation ripples that we measure through time are consistent with the scaling relationship 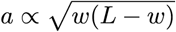 within error (see left panel). The same scatter plot is reproduced in the right panel with uncertainties of ±0.05 cm for each measured length (*a, w* and *L*).

**Figure S6:**
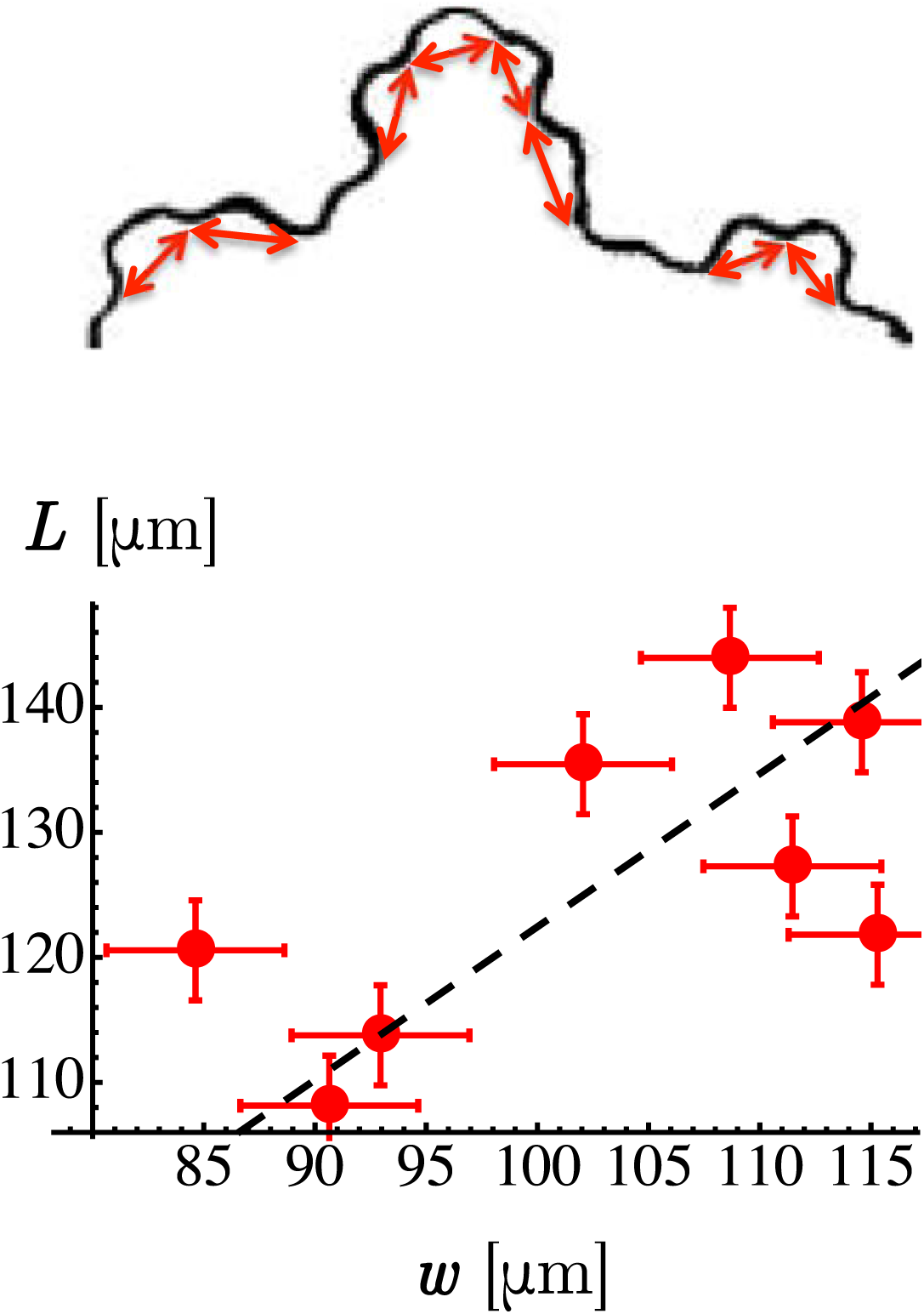
Data from torn plastic sheets. Torn plastic sheets data presented in Fig. 3 of the main text were taken from Fig. 1 in Sharon *et al*., Buckling cascades in free sheets, *Nature* 419:579, 2002. Eight simple ripples were analyzed. They are shown in the top panel of this figure. For these eight ripples, the arc length *L* is plotted as a function of the wavelength *w* in the bottom panel. The slope of the straight line fit (dashed line) is *γ* = 1.22 ± 0.08.

